# Id4 is required for normal ependymal cell development

**DOI:** 10.1101/2021.02.15.431205

**Authors:** Brenda Rocamonde, Vicente Herranz-Pérez, Jose-Manuel Garcia-Verdugo, Emmanuelle Huillard

## Abstract

Ependymal cells are radial glia-derived multiciliated cells lining the lateral ventricles of the brain and spinal cord. Correct development and coordinated cilia beating is essential for proper cerebrospinal fluid flow (CSF) and neurogenesis modulation. Dysfunctions of ependymal cells were associated with transcription factor deregulation. Here we provide evidence that the transcriptional regulator Id4 is involved in ependymal cell development and maturation. We observed that *Id4*-deficient mice display altered ependymal cytoarchitecture, decreased ependymal cell number, altered CSF flow and enlarged ventricles. Our findings open the way for a potential role of Id4 in ependymal cell development and/or motor cilia function.

## 1 Introduction

Ependymal cells are multiciliated epithelial cells organized in a monolayer lining the lateral ventricles (LV) (Doetsch et al., 1997). A subpopulation of radial glia-derived B1 astrocytes present an apical membrane extending a primary cilia that contacts the ventricle (Doetsch et al., 1999; Merkle et al., 2004). Monociliated B1 astrocytes and multicliated ependymal cells are organized within the neurogenic regions of the ventricle wall forming unique pinwheel structures (Mirzadeh et al., 2008).

Ependymal cells are derived from radial glia during embryogenesis between embryonic day 14 (E14) and E16, while maturation occurs later during the first postnatal week. Ependymal cells are born as monociliated epithelial cells (9+0) and their maturation as multiciliated cells (9+2) happens during postnatal day 0 (P0) and P10 (Spassky et al., 2005). It was reported that rotational and translational orientation of basal bodies (BB) are determinant factors of planar cell polarity (PCP) and correlate with CSF flow direction. The coordinated ependymal cell beating is responsible for the correct cerebrospinal fluid (CSF) flow through the ventricular system. CSF flow disruption due to ependymal cell malfunction can lead to hydrocephaly (Taulman et al., 2001) and could impact neuroblast migration towards the olfactory bulb (OB) (Sawamoto et al., 2006).

Factors controlling differentiation and maturation of ependymal cells are not well characterized. It is known that the forkhead transcription factor FOXJ1 is necessary for ependymal cell differentiation from radial glial cells and ciliogenesis (Jacquet et al., 2009). The homeobox factor SIX3 and the transcription factor nuclear factor IX (NFIX) are also involved in ependymal cell development and maturation (Lavado and Oliver, 2011; Vidovic et al., 2018). More recently, it was demonstrated that Geminin and its antagonist GemC1, which are regulators of DNA replication, can determinate the proportion of ependymal cells and neural stem cells (Ortiz-Álvarez et al., 2019). Inhibitor of DNA-binding 4 protein (ID4) is a helix-loop-helix (HLH) protein, acting as a binding partner and modulator of bHLH transcription factors. During embryonic development, ID4 plays an important role in the development of the central nervous system, regulating neural stem cell proliferation and differentiation. ID4-deficient mice present premature differentiation and compromised cell cycle transition of early progenitor cells resulting in smaller brain (Bedford et al., 2005; Yun et al., 2004). However, the role of ID4 in ependymal cell development and maturation from radial glia cells has not yet been addressed. Here we show that ID4 is necessary for correct development of the LV epithelium and for correct ependymal cell maturation. Absence of ID4 during crucial stages of neural fate decision leads to defective ependymal cell development, disrupted planar cell polarity (PCP) and hydrocephalus. Our data suggest for the first time a role for ID4 in ependyma development and function.

## 2 Experimental Procedures

### 2.1 Animals

Mice were housed, bred and treated in an authorized facility (agreement number A751319). All experimental procedures involving mice have been approved by the French Ministry of Research and Higher Education (project authorization number 3572-201601 0817294743 v5). C57BL/6J (Charles River laboratories) and *Id4−/−* (Yun et al., 2004: PMID: 15469968) mice were used at 2-3 months of age. *Glast::CreERT2* (Mori et al., 2006) mice bred to *Id4fl* mice (Best et al., 2014) to obtain *Glast::CreERT2;Id4fl* mice. To induce Id4 deletion, a solution of 60 mg/kg of tamoxifen and 20mg/kg of progesterone diluted in corn oil was administered by oral gavage to pregnant females harvesting embryos at embryonic day 15 (E15). Pups were then obtained at P0 or P10 for wholemount dissection.

### 2.2 Wholemount dissection and immunolabelling

Animals were sacrificed by cervical dislocation. Then, brain was dissected and the whole ventricular-subventricular zone (V-SVZ) was microdissected as described in Mirzadeh *et al.* (Mirzadeh et al., 2010). Fresh tissue was either fixed with 4% paraformaldehyde (PFA; Electron Microscopy Sciences, EMS) and incubated with anti-ZO-1 (1:200, Thermo Fisher Scientific ref. 402200); acetylated tubulin (6-11B, 1:200, Sigma Aldrich ref. T6793) primary antibodies; or fixed with cold 70% ethanol for 10 min and incubated with the anti-gamma tubulin (GTU88, 1:200, Abcam ref. ab11361) primary antibody. Samples were incubated with secondary antibodies Alexa Fluor™ 488 and 596 (1:1000, Life Science Technologies). Then wholemount sections were microdissected and mounted with fluoromount (Sigma, ref F4680). Planar cell polarity (PCP) was determined by the altered orientation of the basal bodies (BB) with respect of the cell wall as described in Mirzadeh *et al*. (Mirzadeh et al., 2010).

### 2.3 Immunofluorescence

Animals were anesthetized with 1 g/Kg sodium pentobarbital (Euthasol) and intracardially perfused with 4% PFA in NaCl 0.9% solution. Brains were dissected and post-fixed in the same solution for 24 h. Fifty micron coronal sections were obtained using a vibratome (Microm). Floating sections were permeabilized with 0.01 M phosphate buffer saline (PBS) containing 0.1% Triton X-100 for 5 min, blocked for 1 hour with 10% Normal Goat Serum (Eurobio Ingen, cat CAECHV00-0U) in PBS–Triton 0.1% (blocking buffer) at room temperature RT and incubated overnight at 4ºC with rabbit anti-ID4 (1:1000, Biocheck ref. BCH-9/82-12) and mouse anti-FOXJ1 (1:500, eBiosc 14-9965-82) antibodies diluted in blocking buffer. Then samples were incubated with anti-mouse Alexa Fluor™ 488 and anti-rabbit Alexa Fluor™ 596 secondary antibodies (1:1000, Life Science Technologies) diluted in blocking buffer. Finally, sections were incubated in DAPI solution for nuclear staining (Invitrogen, cat D3571) and mounted on glass-slides with Fluoromount (Sigma, cat F4680).

### 2.4 Electron microscopy and immunogold staining

For scanning electron microscopy, whole-mount preparations of the lateral wall of lateral ventricles of four animals per group were dissected and fixed with 2 % PFA + 2.5 % glutaraldehyde (EMS) in 0.1 M phosphate buffer (PB) and post-fixed with 1% osmium tetroxide (EMS) in phosphate buffer (PB) for 2 hr, rinsed with deionized water, and dehydrated first in ethanol then with CO_2_ by critical point drying method. The samples were coated with gold/palladium alloy by sputter coating. The surface of the lateral wall was studied under a Hitachi S-4800 scanning electron microscope using Quantax 400 software (Bruker Corporation) for image acquisition.

For pre-embedding immunogold staining, mice were perfused with 4% PFA in 0.1 M PB. Brains were postfixed in in the same fixative solution overnight at 4 °C and sectioned into 50 μm transversal sections using a vibratome. Pre-embedding immunogold staining with rabbit anti-ID4 antibody (1:500; Biocheck) were carried out as previously described (Sirerol-Piquer et al., 2012). Sections were contrasted with 1% osmium tetroxide, 7% glucose in 0.1 M PB and embedded in Durcupan epoxy resin. Subsequently, 1.5 μm semithin sections were prepared, lightly stained with 1% toluidine blue and selected at the light microscope level. Selected levels were cut into 60-80 nm ultrathin sections. These sections were placed on Formvar-coated single-slot copper grids (Electron Microscopy Sciences) stained with lead citrate and examined at 80 kV on a FEI Tecnai G^2^ Spirit (FEI Company) transmission electron microscope equipped with a Morada CCD digital camera (Olympus).

### 2.5 Image acquisition and analysis

Fluorescent images were obtained using a Zeiss ApoTome 2 Microscope or Olympus Confocal microscope FV1000. Images were analysed using ImageJ software.

### 2.6 Statistical analysis

Statistical analysis was performed using GraphPad Prism 6. Unless otherwise indicated in the figure legends, non-parametric Mann-Whitney test was used to compare experimental and control groups. Values are expressed as mean ± standard deviation.

## 3 Results

### 3.1 ID4 is expressed in ependymal cells from the LV

We performed ID4 immunolabelling on wholemount preparations of the LV of adult C57BL/6J mice (Figure 1A). We detected the expression of ID4 protein in ependymal cells as can be observed by co-localization with FOXJ1 protein (Figure 1B). To confirm this observation, we performed immuno-gold labelling for the ID4 protein in ultrathin sections from the V-SVZ. Several cell types, such as B1 astrocytes and progenitor stem cells were positive for ID4-immungold. In addition, ID4 labelling was detected in ependymal cells lining the LV confirming our immunofluorescence results (arrowheads in Figure 1C).

**Figure 1.**
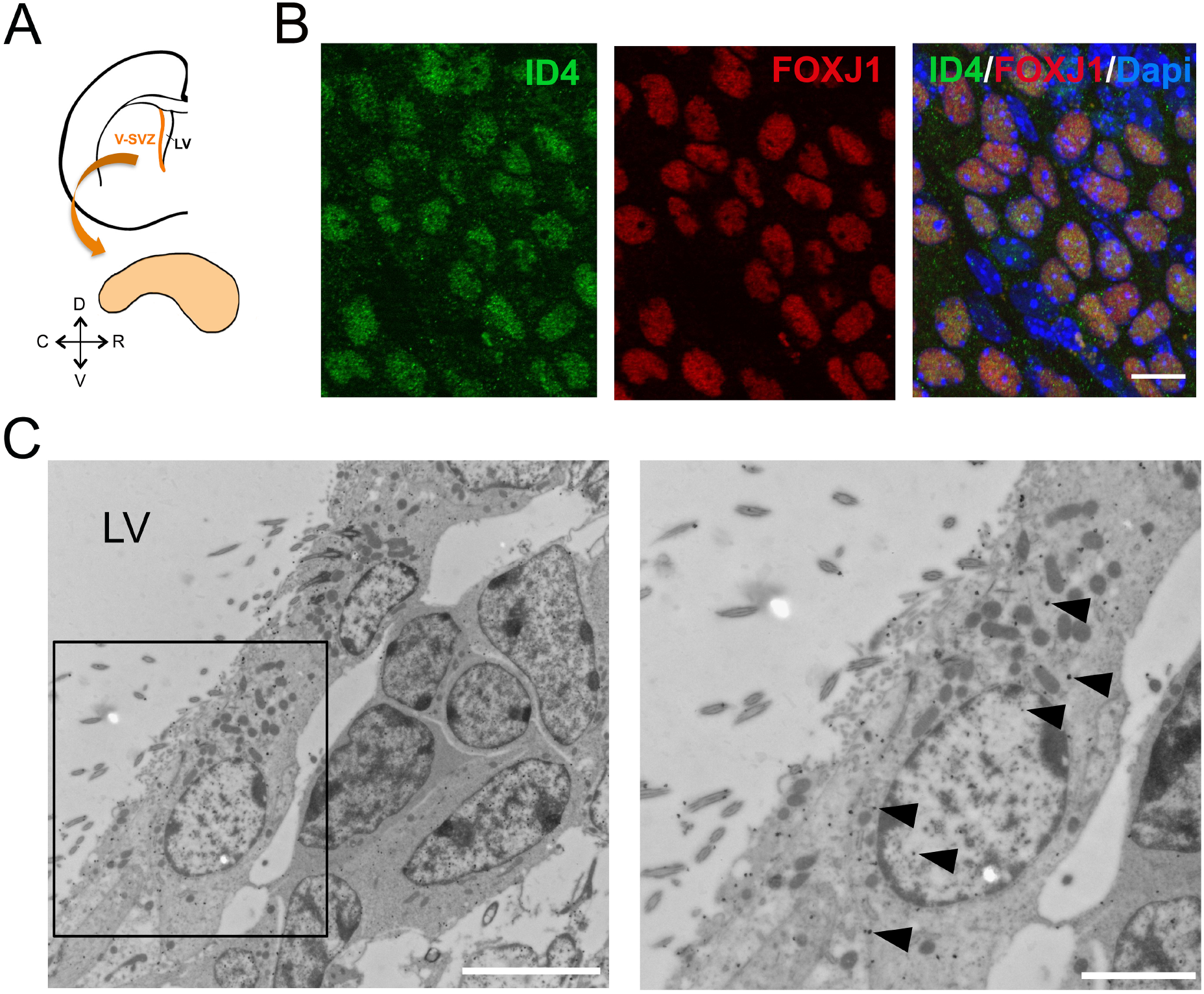
ID4 expression in ependymal cells. A) Scheme of a coronal section of a mouse brain hemisphere showing the ventricular-subventricular zone (V-SVZ) of the lateral ventricles (LV) and a wholemount section that can be dissected out from this region. B) Immunolabelling of ID4 and FOXJ1 (ependymal cell marker) on wholemount sections of adult C57BL/6J mice. Scale bar: 5 μm. C) Immuno-gold staining of ID4 protein on V-SVZ sections. Left panel shows ependymal cells next to astrocytes and neuroblasts. Arrowheads indicate some of the gold particles labelling Id4 protein in ependymal cells. Right panel shows higher magnification of boxed area on left panel. Scale bar: 5 μm left and 2 μm right.

### 3.2 *Id4−/−* mice display defects in LV development and ependymal cells function

In order to investigate the role of ID4 in ependymal cells, we analysed the V-SVZ of *Id4−/−* mice (Id4KO) (Yun et al., 2004). Id4KO mice consistently displayed enlarged ventricles (Figure 2A-B). This was associated with thinning of the ventricular wall and stretching of the ependymal cells, as observed by light microscopy in toluidine blue-stained semithin sections (Figure 2C). Scanning electron microscopy of wholemount preparations revealed the absence of adhesion point ‒the area of the lateral and medial ventricle wall that adheres to each other– in *Id4−/−* mice (Figure 2D). In addition, we observed that ependymal cell density was decreased in all three rostral, central and caudal areas of the ventricle wall.

**Figure 2.**
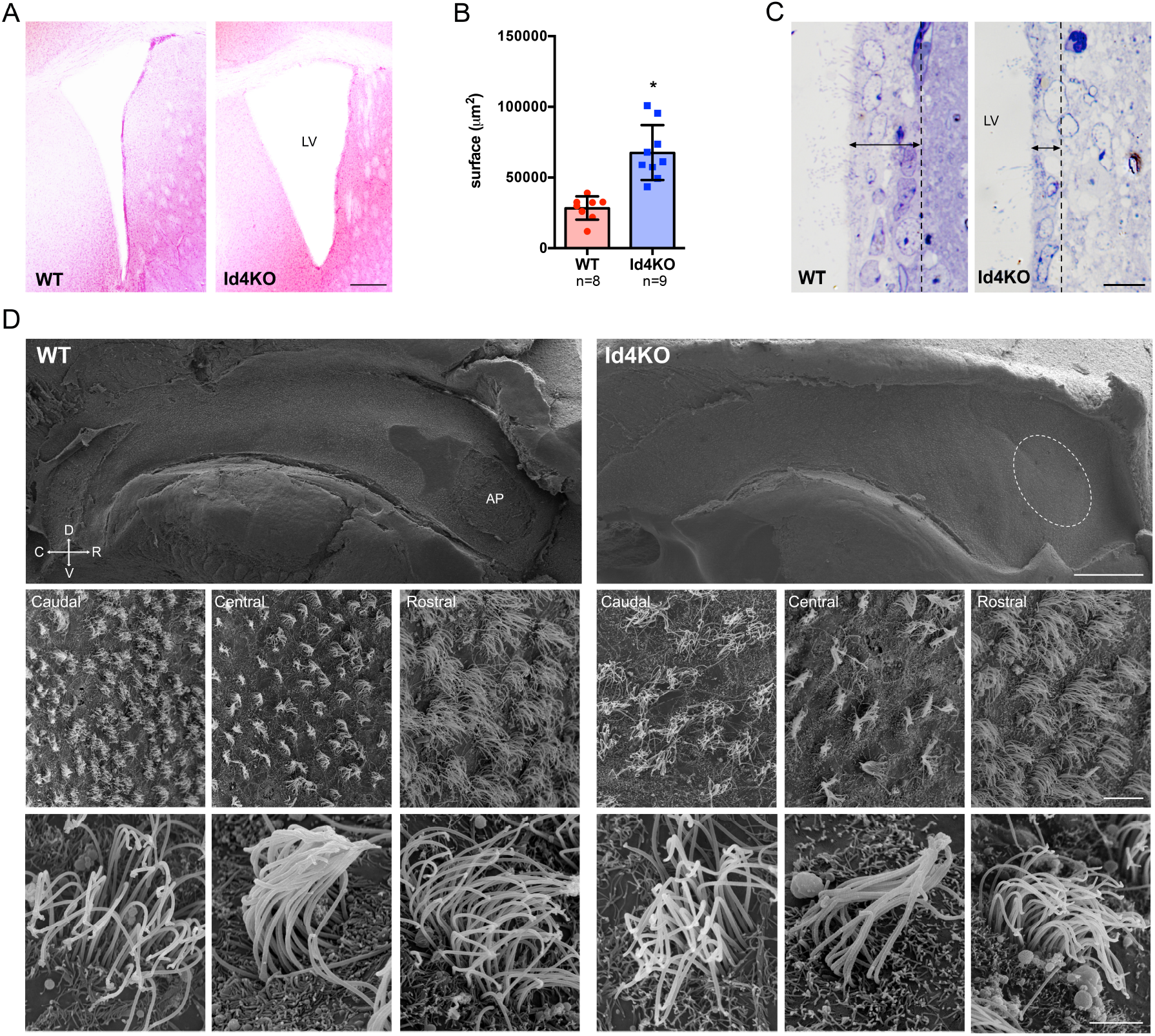
Absence of ID4 during development results in enlarged ventricles. A) Haematoxylin-eosin staining of brain coronal sections from *Id4+/+* (WT) and *Id4−/−* (Id4KO) adult mice showing enlarged lateral ventricles (LV). Scale bar: 200 μm. B) Quantification of the surface of the LV in WT and Id4KO mice. *p-value ≥ 0.05. C) Semithin sections of WT and Id4KO mice present a thinner subventricular wall. Scale bar: 5 μm. D) Scanning electron microscopy of wholemount preparations from WT and Id4KO mice show enlargement of the ventricles and disappearance of the adhesion point (AP), decreased ependymal cell density and altered organization in cilia. Scale bar from up to down: 500 μm, 5 μm and 2 μm.

To confirm a decrease in ependymal cell density, we performed immunofluorescent staining on wholemount preparations of the cell wall (labelling tight junctions with ZO-1 antibody) and cilia (acetylated tubulin, 6-11B) in WT and Id4KO mice (Figure 3A). The density of ependymal cells was significantly decreased in Id4KO mice together with an increase in the cell surface when compared the same regions (Figure 3B-C). To investigate whether altered ependyma in Id4KO brains might lead to altered CSF flow, we evaluated planar cell polarity of ependymal cells (PCP) by measuring cilia basal bodies (BB) orientation (Mirzadeh et al., 2010). Basal bodies were labelled with anti-γ-tubulin (GTU88) antibody and its orientation was determined relative to the ependymal cell wall labelled anti-ZO1 antibody. We noticed that the organization of BB patches was altered in the Id4KO mice, with a significant decrease of the median orientation (Figure 3D,E).

**Figure 3.**
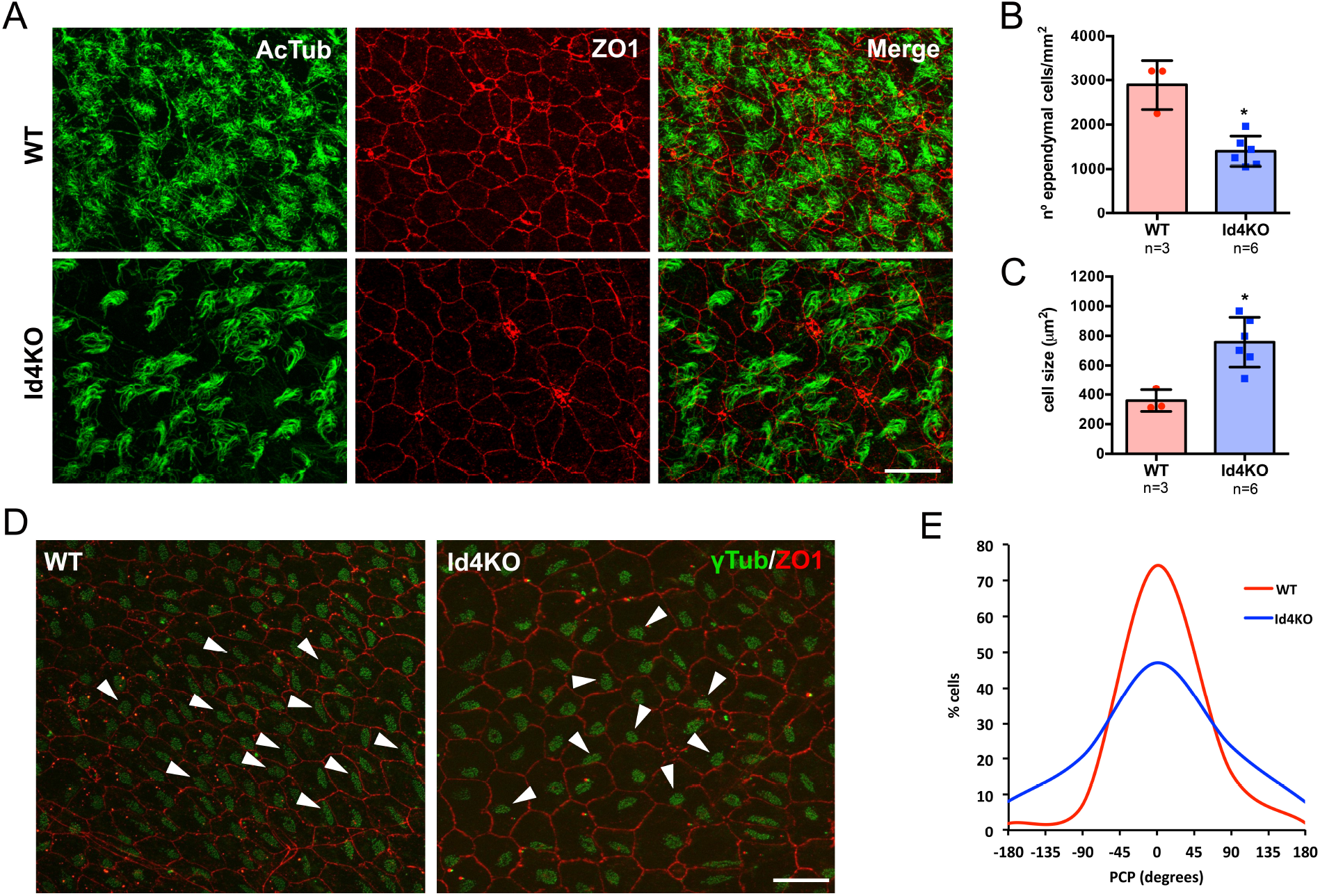
ID4KO mice present disrupted PCP. A) Immunofluorescence for ZO1 (red) and acetylated tubulin (green) in wholemount preparations of WT and Id4KO mice. Scale bar: 10 μm. B) Quantification of the number of ependymal cells determined by ZO1 staining. C) Quantification of ependymal cell ventricular surface. D) Immunofluorescence for ZO1 (red) and γ-tubulin (green) in wholemount preparations show disorganized planar cell polarity (PCP). Scale bar: 10 μm. E) Quantification of the PCP in the wholemount preparations of WT and Id4KO mice. \**p*-value ≤ 0.05.

### 3.3 Id4 deletion during embryogenesis impacts on ependymal cell development

Ependymal cells differentiate from radial glia cells at E14-16 and maturation occurs from caudal to rostral orientation during the first postnatal weeks (Spassky et al., 2005), where the primary cilium is replaced by multiple motile cilia (9+2) (Mirzadeh et al., 2010). To evaluate whether ID4 was involved in ependymal cell differentiation and/or maturation we induced Id4 deletion in *GlastCreERT2-Id4flox (Id4cKO)* mice at E15 and analysed their phenotype at P0 to evaluate differentiation and at P10 to analyse maturation (Figure 4A). We performed immunofluorescence for γ-tubulin (GTU88) to identify the BB and for Z01. The defects in ependymal cell maturation were already present at P0 but were more evident at P10 (Figure 4B). In addition, the number of ependymal cells seemed to be decreased already at P0, suggesting that Id4 may be involved in differentiation of ependymal cells at embryonic stages. Quantification of the number of ependymal cells showed a significant decrease in Id4cKO in rostral regions at P0 (Figure 4C). In addition, evaluation of the presence of matured ependymal cells at P10 showed a decline in Id4cKO LV, although it did not reach statistical significance. Together, our data suggest that Id4 may be important for ependymal cell maturation and correct cilia development.

**Figure 4.**
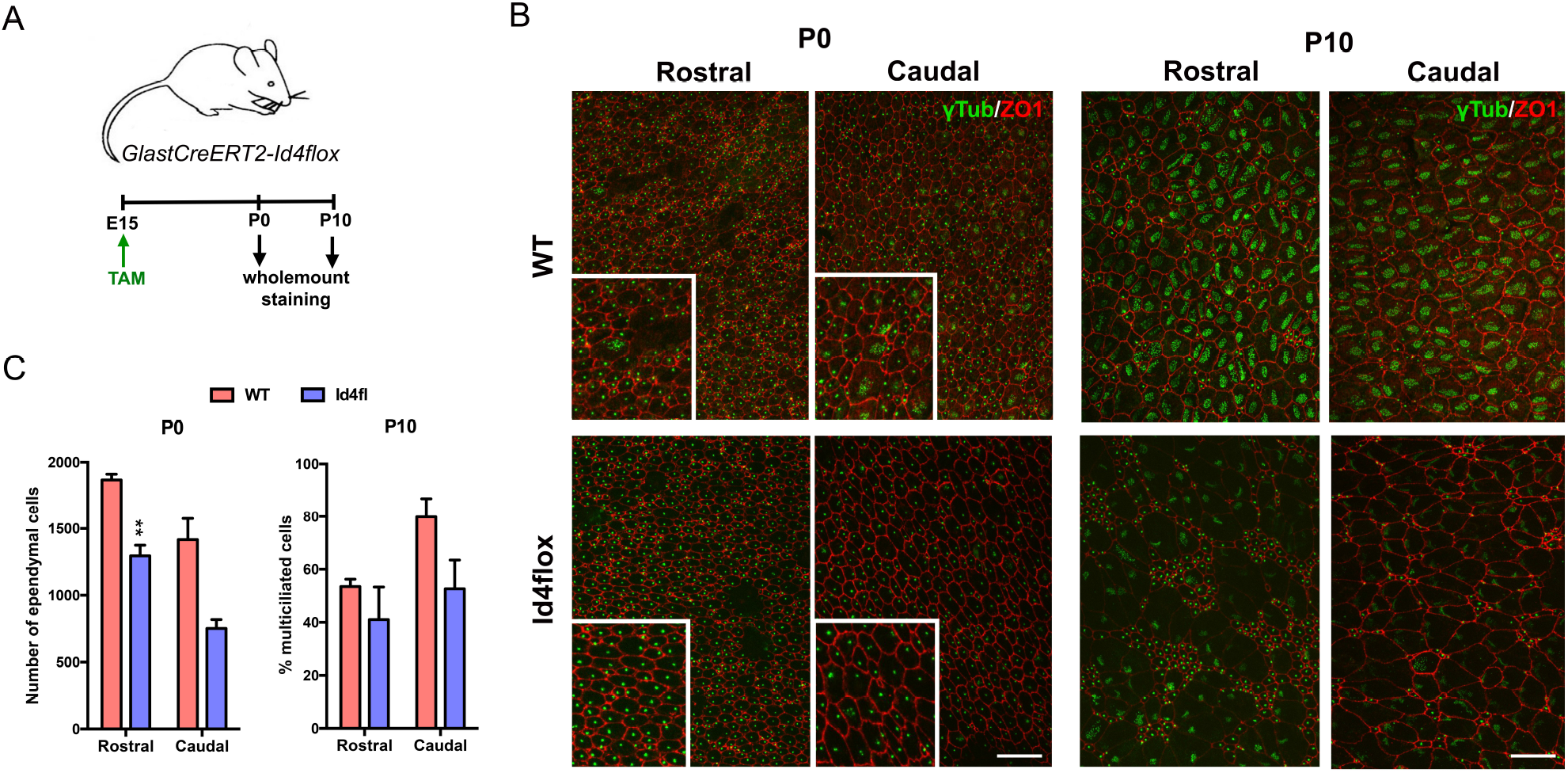
Absence of ID4 results in delayed ependymal cell maturation. A) Scheme of the experimental set-up. B) Wholemount staining of P0 and P10 mice for Z01 (red) and γ-tubulin (green) showing defective ependymal cell development. Scale bar: 10 μm. C) Number of ependymal cells at P0 and the frequency of differentiated ependymal cells (% of multiciliated cells) at the time of the analysis in rostral and caudal regions of the lateral ventricle.

## 4 Discussion

ID4 plays an essential role in correct neural cell differentiation and maturation during embryogenesis (Yun et al., 2004; Bedford et al., 2005). In this work, we present evidence that ID4 might also be important for development and maturation of ependymal cells from radial glial cells. First, we describe for the first time that ID4 protein is expressed in ependymal cells. Absence of ID4, initiated at early stages of brain development, resulted in altered ependymal cell layer cytoarchitecture, decreased ependymal cell numbers and enlarged ventricles as a consequence. The absence of adhesion point was also a constant in Id4KO mice and was already reported to be linked to hydrocephalus in other mutants, such as KIF3A-deficient mice (Mirzadeh et al., 2010). Early inactivation of *Id4* at E15 during differentiation of ependymal cells from radial glial cells resulted in decreased number of ependymal cells, suggesting that ID4 may be important in ependymal cell determination from radial glial cells. Defects in differentiation from radial glial cells were also observed when the forkhead transcription factor FOXJ1 was absent (Jacquet et al., 2009). GemC1 – a regulator of the DNA replication– was another factor playing an important role in the ependymal cell-neural stem cell balance (Ortiz-Álvarez et al., 2019).

In addition to a defective differentiation, our findings suggest that ID4 may also be involved in ciliogenesis. Delayed ciliogenesis was also detected in ependymal cells at P10 when Id4 was deleted. Another transcription factors involved in ciliogenesis were FOXJ1 and NFIX (Jacquet et al., 2009; Vidovic et al., 2018). Decreased number and delayed maturation of ependymal cell beating capacity could lead to disrupted CSF flow dynamics early during brain development and accumulation of CSF. Accumulation of CSF within the brain ventricles due to defect in ependymal cell development is one of the mechanisms responsible of hydrocephalus (Ibañez-Tallon et al., 2004). Despite complete cilia development in ependymal cells, PCP seemed to be affected by the absence of Id4. This could suggest a potential role of Id4 in centrosome organization or cilia motility. Malfunction in ependymal cell beating activity can impair CSF clearance and cause excessive accumulation within the ventricles leading to hydrocephalus as a result of increased pressure. On the other hand, CSF flow dynamics can also impact neural stem cell (NSC) proliferation and neuroblast migration towards the olfactory bulb. Several studies have reported that the NSC’s primary cilium works as an “antenna” sensing the changes in CSF flow (Silva-Vargas et al., 2016). CSF flow generates protein gradients contributing with vector information for migratory neuroblasts (Sawamoto et al., 2006). Therefore, disruption of CSF caused by genetic defects could also indirectly impact NSC homeostasis and/or migration.

Our findings here show for the first time a role of Id4 in ependymal cell differentiation and maturation. Further investigation should be conducted to better understand the mechanism leading to such phenotype, the potential protein partners of Id4 and the impact on neurogenesis and neuroblast migration.

## 6 Author Contributions

Conceptualization: B.R. and E.H.

Methodology: B.R. and E.H.

Investigation: B.R., V.H-P., V.S-V.

Writing – Original Draft: B.R. and E.H.

Writing – Review & Editing: all authors

Funding Acquisition: J-M.G-V., V.H-P., B.R. and E.H.

Supervision: J-M.G-V. and E.H.

## 8 Acknowledgements

We thank Mark Israel, Jane Visvader and Magdalena Götz for providing Id4KO, Id4fl and GlastCreERT2 mice, respectively. We thank Nathalie Spassky for reading and commenting the manuscript.

